# Theta bursts precede, and spindles follow, cortical and thalamic downstates in human NREM sleep

**DOI:** 10.1101/260786

**Authors:** Chris Gonzalez, Rachel Mak-McCully, Burke Rosen, Sydney S. Cash, Patrick Chauvel, Hélène Bastuji, Marc Rey, Eric Halgren

**Affiliations:** Neurosciences, University of California San Diego, La Jolla, California 92093; University California Berkeley, Berkeley, California 94720; Department of Neurology, Massachusetts General Hospital and Harvard Medical School, Harvard University, Boston, Massachusetts 02114; Aix-Marseille Université, Marseille 13385, France; Central Integration of Pain, Lyon Neuroscience Research Center, INSERM, U1028; CNRS, UMR5292; Université Claude Bernard, Lyon, Bron, France; Departments of Radiology and Neurosciences, University of California, San Diego, California 92093

**Author notes:** indicates primary authors Corresponding author: Chris Gonzalez, Neurosciences, University of California San Diego, La Jolla, California 92093. **Financial disclosures:** The authors declare no competing financial interests or conflicts of interests. **Abbreviated Title**: Theta precedes downstates, spindles follow.

## Abstract

Since their discovery, slow oscillations have been observed to group spindles during non-REM sleep. Previous studies assert that the slow oscillation downstate (DS) is preceded by slow spindles (10-12Hz), and followed by fast spindles (12-16Hz). Here, using both direct transcortical recordings in patients with intractable epilepsy (n=10, 8 female), as well as scalp EEG recordings from a healthy cohort (n=3, 1 female), we find in multiple cortical areas that both slow and fast spindles follow the DS. Although discrete oscillations do precede DSs, they are theta bursts (TB) centered at 5-8Hz. TBs were more pronounced for DSs in NREM stage N2 compared with N3. TB with similar properties occur in the thalamus, but unlike spindles they have no clear temporal relationship with cortical TB. These differences in corticothalamic dynamics, as well as differences between spindles and theta in coupling high frequency content, are consistent with NREM theta having separate generative mechanisms from spindles. The final inhibitory cycle of the TB coincides with the DS peak, suggesting that in N2, TB may help trigger the DS. Since the transition to N1 is marked by the appearance of theta, and the transition to N2 by the appearance of DS and thus spindles, a role of TB in triggering DS could help explain the sequence of electrophysiological events characterizing sleep. Finally, the coordinated appearance of spindles and DSs are implicated in memory consolidation processes, and the current findings redefine their temporal coupling with theta during NREM sleep.

**Significance Statement:** Sleep is characterized by large slow waves which modulate brain activity. Prominent among these are ‘downstates,’ periods of a few tenths of a second when most cells stop firing, and ‘spindles,’ oscillations at about twelve times a second lasting for about a second. In this study, we provide the first detailed description of another kind of sleep wave: ‘theta bursts,’ a brief oscillation at about six cycles per second. We show, recording during natural sleep directly from the human cortex and thalamus, as well as on the human scalp, that theta bursts precede, and spindles follow downstates. Theta bursts may help trigger downstates in some circumstances, and organize cortical and thalamic activity so that memories can be consolidated during sleep.

## Introduction

During NREM sleep, the brain endogenously produces electrical activity dominated by larger amplitude, lower frequency (0.1-16 Hz) rhythms compared with wake or REM states. Downstates (periods of neuronal quiescence lasting a few hundred ms) and sleep spindles (approximately 0.5-2s, 10-16 Hz oscillations) are two canonical NREM events with initiating mechanisms largely attributed to cortical and thalamic activities, respectively. However, the extensive bidirectional connections between cortex and thalamus precludes a simple entraining mechanism for either event, and properties such as duration (Bonjean et al. 2011; Bazhenov et al. 2002; Bartho et al. 2014), frequency of occurrence (Timofeev et al. 2000), and synchrony (Contreras et al. 1996) are all shaped by their cooperative dynamics (Crunelli & Hughes 2010; Steriade 1997). These rhythms are believed to serve functional roles in sleep-dependent memory consolidation (Sejnowski & Destexhe 2000; Diekelmann & Born 2010; Hanert et al., 2017). In particular, the specific grouping of spindles by slow waves has been associated with improved declarative memory in humans (Niknazar et al. 2015; Mölle et al. 2011) and fear conditioning in mice (Latchoumane et al. 2017).

While several studies report faster spindle frequency activity (>12 Hz) occurs on the transition from the down to up state (Mölle et al. 2011; Cox et al. 2014; Andrillon et al. 2011; Klinzing et al. 2016), the temporal relation of lower frequency (4-12 Hz) content to DSs has not been definitively established. This relationship is more clear during NREM stage N2 sleep when DS usually occur without preceding upstates as the main component of the K-complex (KC) (Cash et al. 2009). Some authors observe “polyphasic waves […]just before the onset of the negative K-complex sharp wave” (Rodenbeck et al. 2006), or an approximately 7 Hz short-lasting “intra-KC” oscillation (Kokkinos & Kostopoulos 2011; Kokkinos et al. 2013). More recently, however, the upstate to DS transition has been associated with slow spindle activity (9-12 Hz) (Mölle et al. 2011; Klinzing et al. 2016; Yordanova et al. 2017). Findings from the latter studies bolster the hypothesis that there are distinct types of spindles, slow and fast, which could have their own rhythmogenesis mechanisms (Timofeev & Chauvette 2013; Fogerson & Huguenard 2016). Here, we present findings that challenge this hypothesis using bipolar transcortical and thalamic recordings from epileptic patients to obtain focal measures of DSs and spindles. We confirm these findings at the scalp, using EEG recordings from a non-clinical population. We propose that regardless of frequency, spindles recorded either intracranially or at the scalp, are more likely to start on the down to up transition, and we describe a theta burst distinct from spindles that can accompany the up to down transition, especially in stage N2 NREM sleep.

## Materials and Methods

### Intracranial Recordings

Stereoencephalography (SEEG) was obtained in 10 patients (8 female; mean ±std. dev. age: 38.7 ±12.7) undergoing evaluation for pharmaco-resistant epilepsy at Massachusetts General Hospital, La Timone Hospital, Marseille, France, or Neurological Hospital, Lyon, France. At Massachusetts General Hospital, electrode contacts were localized using computed tomography (CT) of the implanted electrodes superimposed on preoperative MRI (Dykstra et al. 2012). Each SEEG electrode had either 8 (5mm center-to-center spacing) or 6 contacts (8mm spacing). Each contact was 1.28 mm in diameter and 2.4mm long. Signals were sampled at 500 Hz and band pass filtered from 0.33 to 128 Hz. At La Timone Hospital, localization of electrode contacts was performed using MRI and CT of implanted electrodes. For one patient, localization was determined using preoperative MRI and surgical planning. Each electrode had either 10 or 15 contacts (3.5mm center-to-center spacing). Each contact was 0.8mm in diameter and 2mm long. The recordings were sampled at 256, 512, or 1024 Hz. At the Neurological Hospital, electrode localization was determined directly from stereotactic teleradiographs without parallax performed within the stereotactic frame (Talairach & Tournoux 1998). These locations were superimposed onto the preimplantation 3T structural MRI (3D MPRAGE T1 sequence) after alignment with the skull. The locations of cortical and thalamic contacts were determined by reference to the atlases of Duvernoy (H Duvernoy 1999) and Morel et al (Morel et al. 1997). Each electrode had either 10 or 15 contacts (3.5 mm center-to-center spacing). Each contact was 0.8mm in diameter and 2mm long. The recordings were sampled at 256 Hz and band pass filtered from 0.33 to 128 Hz. Informed consent was obtained from all patients.

SEEG recordings were bipolar between adjacent contacts spanning the cortical ribbon. These ‘transcortical bipolar contacts’ provide relatively focal measurements of the local field potentials generated in the transected cortex (Mak-McCully et al. 2015). The polarity of bipolar derivations was inverted if necessary to assure that DS (as confirmed with decreased HG- see below) were negative. This produced recordings with a relatively consistent relationship to the underlying cortical generators, as evidenced by the consistent phase relation between HG and TB (see below). They were chosen as previously described (Mak-McCully et al. 2015). Only channels with both slow oscillations and spindles apparent to visual inspection were included for analysis. Channels were excluded if a clinical electroencephalographer judged significant interictal activity, pathological background changes, or early involvement in the ictal discharge. At each site, sleep scoring was performed by clinical experts (SC, HB, MR) using examination of bipolar electrodes (N=7) or scalp EEG with EOG/EMG when available (N=3). Analyses were performed on N2 and N3 sleep periods only (Silber et al. 2007).

In total, 60 cortical bipolar SEEG channels obtained from 10 patients and 8 thalamic (mainly pulvinar) bipolar channels from three of the 10 patients during NREM sleep were included for analysis. The locations of all cortical bipolar channels are shown in Figure 1A, as well as examples of bipolar derivations for cortical (Fig. 1B) and thalamic (Fig. 1C) sites. On average, each patient had 6 cortical channels (ranging from 2 to 13) and 120.6 minutes of NREM sleep from one night (ranging from 26.5 to 240 minutes). Downstates and spindles were detected on each channel separately using previously published methods (see below) (Mak-McCully et al. 2017).

**Figure 1.**
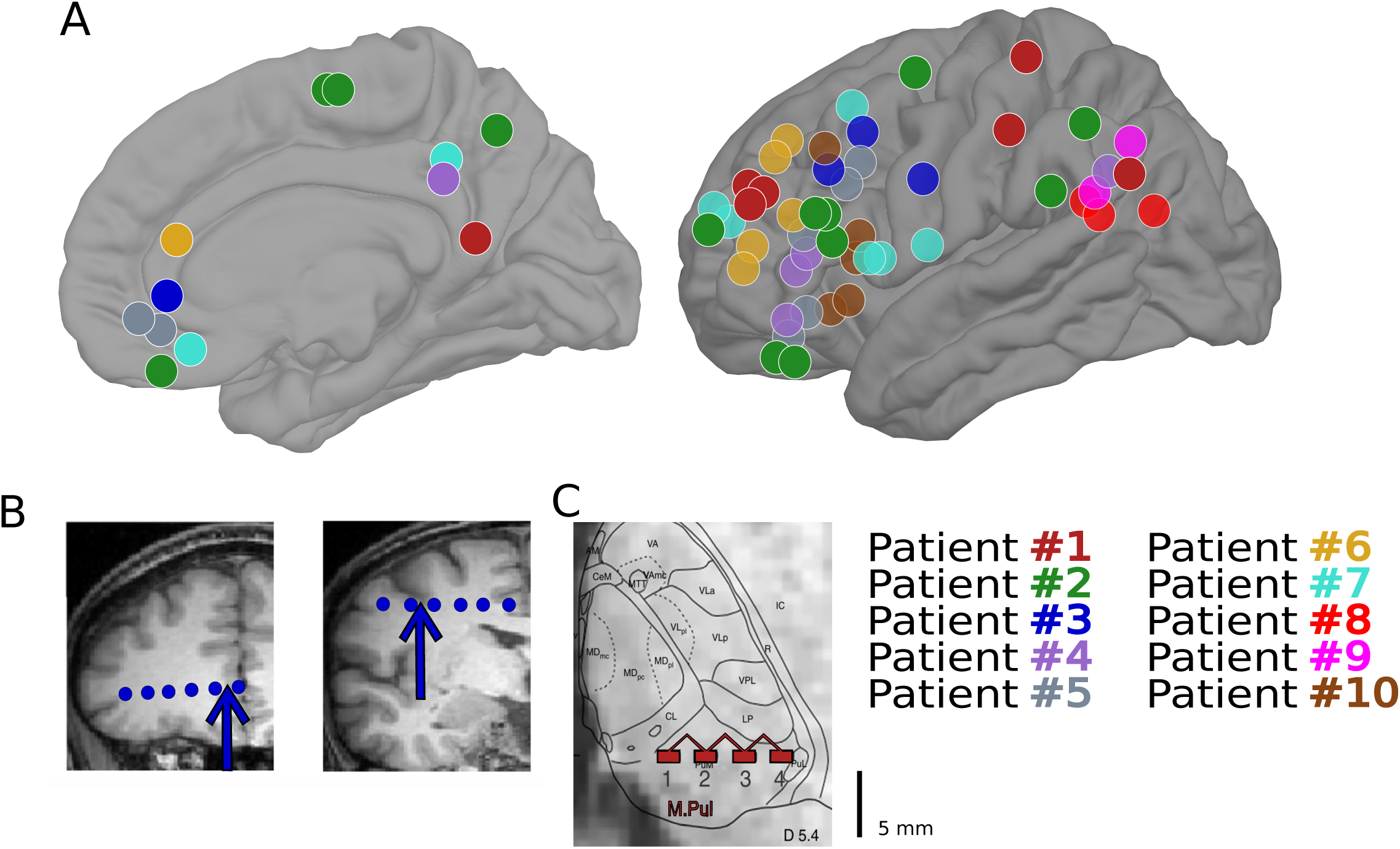
Bipolar SEEG recordings obtained from epileptic patients. A) Locations of 60 cortical bipolar recordings from 10 patients with intracranial electrodes. B) Illustration of bipolar transcortical derivation used throughout the study, with arrows indicating two cortical bipolar channels from Patient 3. C) Illustration of bipolar recordings from Patient 1 from the Pulvinar of the thalamus.

### Scalp EEG Recordings

Scalp EEG data were recorded during sleep from 3 healthy subjects (1 female). Written informed consent approved by the Partners Healthcare Network was obtained for all subjects prior to their participation. Subjects wore a 70 channel EEG cap with a modified 10-20 montage (Elekta Neuromag). Data were referenced to left mastoid. Magnetoencephalographic data were collected simultaneously but are not reported here. Periods of N2 sleep were identified according to standard criteria (Iber et al. 2007). Gross artifacts were removed by visual inspection.

### Downstate detection

Downstates were detected on each channel as follows: 1) Apply a zero-phase eighth order Butterworth filter from 0.1 to 4Hz; 2) Select consecutive zero crossings within 0.25 to 3 seconds; 3) Calculate amplitude peak between zero crossings and retain only the bottom 20% of peaks for intracranial recordings, or the bottom 10% of peaks for scalp EEG recordings.

For downstate detection in intracranial recordings, only periods of N2 and N3 sleep free of visually-identified epileptiform discharges were used. Bipolar SEEG channels exhibiting downstates were also required to show decreases in power within 60-100Hz (High Gamma; HG) exceeding 1 dB within ± 250 ms of the negative DS peak. Sixty such channels were identified, showing mean decreases in HG power during DS troughs of -3.18 dB (range -1 to -8). Downstate masks were padded with ± 250ms centered around the trough when determining the extent cortical events overlapped with thalamic events.

### Spindle detection

The current clinical standard for sleep scoring adopted by the American Academy of Sleep Medicine is 11-16 Hz (Silber et al. 2007), but the major previous publications describing sleep spindles using intracranial recordings in humans adopted 9-16 Hz (Andrillon et al. 2011; Piantoni et al. 2017) or 10-16 Hz (Mak-McCully et al. 2017; Hagler et al. 2018). Recent analyses with scalp recordings of fast vs slow spindles have also used either 9-15 Hz (Mölle et al. 2011; Klinzing et al. 2016) or 10-16 Hz (Cox et al. 2014) or even 9-16 Hz (Yordanova et al. 2017), but in all cases with the division between fast and slow spindles at 12 Hz. Here we consider spindles as 10-16 Hz events and define spindles ≤12 Hz as slow. Recordings were notch filtered (either 49 to 51 Hz or 59 to 61 Hz, depending on country of origin) and then band passed at 10-16 Hz using a zero-phase frequency domain filter (transition bands 30% of cutoff frequency). Taking the absolute value of this filtered signal produced a spindle-band amplitude envelope. This envelope was convolved with a 400ms Tukey window, the median amplitude is subtracted, and normalized by the median absolute deviation. This signal was used to detect the onset and offset of putative spindle epochs. To detect the middle of spindle epochs, we convolved the amplitude signal with a 600ms Tukey window, normalized as before, and identified peaks with magnitude larger than 2 for intracranial recordings and larger than 1 for scalp EEG recordings. Then, we defined the onset and offset as 40% of the peak amplitude of the original spindle amplitude envelope. Any overlapping or duplicate epochs were resolved, and epochs less than 300ms excluded. We then applied a series of strict exclusion criteria for putative spindle epochs. These included removing any epochs that also exceeded 5 for a low (4-8Hz) or high (18-25 Hz) amplitude envelope. We also required 5 peaks in a broad-band filtered signal (4-25 Hz) with an amplitude greater than the median absolute deviation per channel and at least 25% amplitude of the largest peak. Spindles in the thalamus were detected using a modified version of this detector as previously reported (Mak-McCully et al. 2017).

### Theta burst detection

Theta bursts were detected by modifying a previously reported spindle detector (Andrillon et al. 2011). Our procedure was as follows: (1) Apply a zero-phase eighth order Butterworth filter from 5-8Hz (range selected to minimize overlap with delta and spindle content); (2) Calculate the mean of the Hilbert envelope of this signal smoothed with a Gaussian kernel (300ms window; 40ms sigma); (3) Detect events with +3SD threshold for the peak and identify the start and stop times with a +1SD threshold; (4) Only include events with a duration between 400ms and 1s; and (5) for each band pass-filtered peak in putative burst, calculate the preceding trough-to-peak deflection, and only take events that have at least 3 peaks exceeding 25% of the maximum deflection. We also required bipolar SEEG recordings to have at least 50 TBs before calculating average theta frequency or the proportion of theta events associated with DSs, which excluded four of sixty cortical channels.

### Phase-amplitude coupling (PAC)

For the intracranial recordings, we correlated the phase of either spindle (10-16 Hz) or theta (5-8 Hz) with the analytic amplitude in 60-100 Hz for all cortical channels across subjects. We chose 60-100Hz to keep the range consistent across subjects, and because our lowest sampling frequency was 256 Hz. Only channels that had at least 30 theta events and 30 spindles were included (58 of 60 cortical channels). We band passed our data in the theta, spindle, and gamma frequency ranges using finite impulse response filters with an order equal to the duration of three cycles of the lowest frequency. We used the Hilbert transform to extract the analytic signal from our band passed data, took the phase angle from our theta and spindle band passed data, and the amplitude of our HG band pass signal. Correlations between the phase of the lower frequency signal and HG amplitude were evaluated across a duration equal to the first two cycles of the lower frequency range per signal, across all detected events, for each channel. The observed phase-amplitude coupling (PAC) measure was calculated by taking the length of the average complex vector of the low-frequency phase, weighted by the corresponding high-frequency power time series (Canolty et al. 2006). Significance was assessed for each channel and event type using non-parametric permutation statistics. Specifically, for each channel and event type, the phase time series was randomly offset relative to the power time series and PAC re-calculated 1000 times, generating a null distribution to compare our observed PAC measure against. The preferred phase was determined only for channels with a PAC-Z value greater than 3 (p<0.002).

### Time domain and spectral analyses

We created event-related histograms to quantify the timing of intracranially-recorded spindle or theta events relative to all intracranially-recorded DS troughs. For each bipolar SEEG recording, we required at least 30 events associated with a DS within ±1 s of the DS trough to be included in the grand average histograms. Furthermore, we required at least 20 events within +/- 500ms to assess if spindle (theta) events were more likely to start 500ms after (before) the DS trough using a two-way binomial test. Binomial tests were corrected for multiple comparisons using Bonferroni correction (alpha= 0.05) for each graphoelement type. Data processing and analysis was performed in MATLAB, and time-frequency plots created using EEGLAB (Delorme & Makeig 2004).

We also examined the spectral profile of periods prior to downstates. Power-spectral densities (PSDs) were computed for the epoch 500 ms to 0 ms before the DS trough. For each channel, these were ranked and quartiled by the power in the 4-12 Hz band. PSDs in the first and fourth quartile were normalized as z-scores, and averaged over downstates. The difference between the quartiles is plotted in Figures 2G (SEEG) and 5E (scalp EEG).

**Figure 2.**
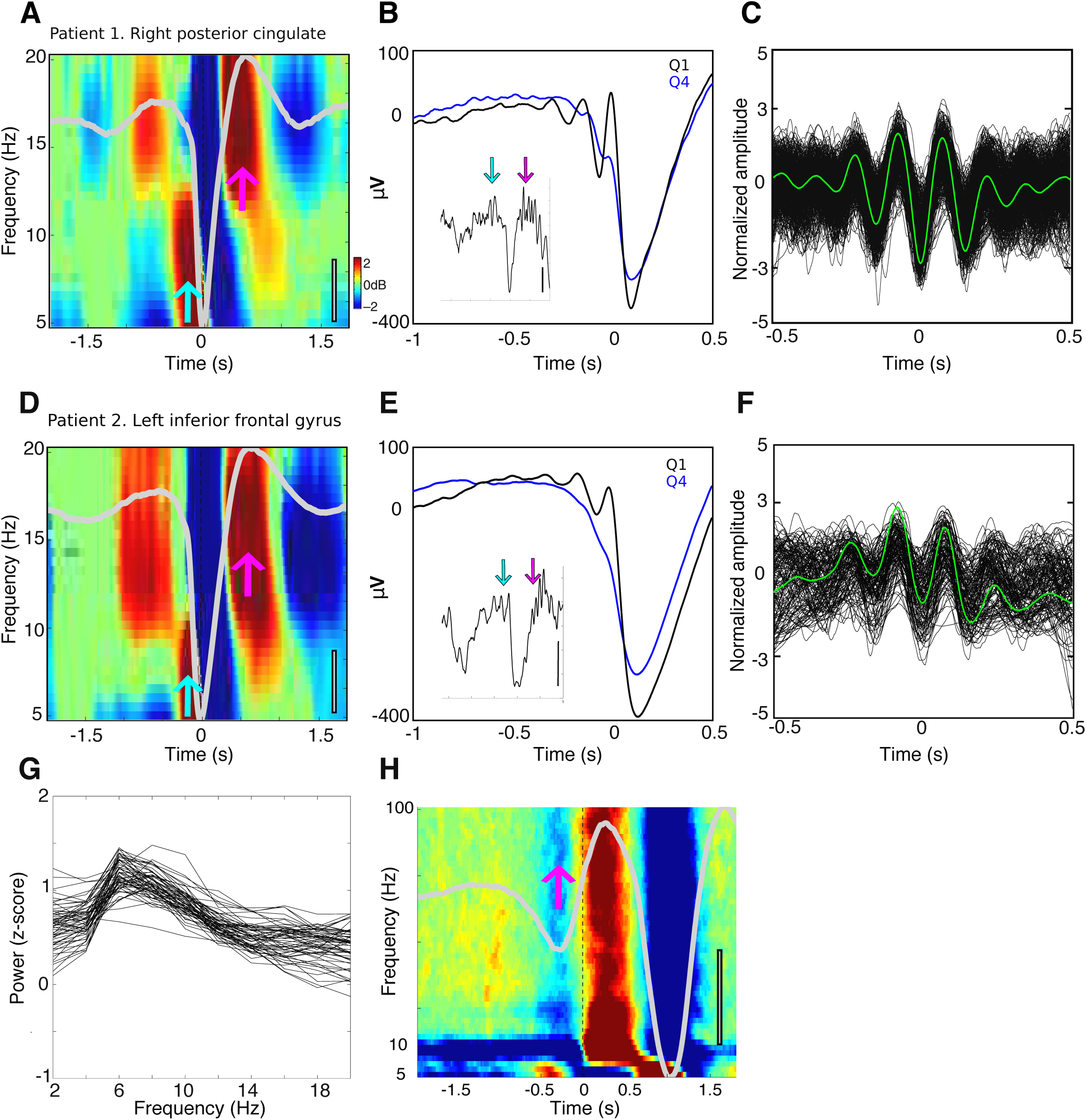
Cortical theta during NREM sleep. A-C) Patient 1, right posterior cingulate. D-F) Patient 2, left inferior frontal gyrus. For both A (n=2084) and D (n=1265), time-frequency plots locked to DS troughs reveal increases in the theta range (cyan arrows) immediately before the trough, and centered within spindle range (pink arrows) from 0.5 to 1 s after the trough (and more weakly from -1 to -.5 s prior to the trough, teal arrows, due to a preceding DS-see panel H). The entire epoch was used as baseline. The average of all DSs is overlaid in light gray, gray scale bars indicate 100µV. B,E) Average, unfiltered LFP time-locked to the first filtered positive theta peak prior to the DS trough. Black indicates the top quartile (n=521 for B, n=316 for E), blue the bottom quartile of DSs with prior theta power. Insets reveal a single trace from the top quartile (Scale bars indicate 200µV for inset, time ranges from -2 to +.5 s of the DS trough). Cyan and pink arrows indicate theta events and spindles, respectively. Positive potentials indicate cortical surface positivity. C,F) Raw LFP of detected theta events locked to the deepest trough, z-normalized. Average shown in green for C (n=509) and F (n=148). G) Pre-downstate spectra. Plotted are the difference of the first and fourth quartile average power spectral densities of 500 ms pre-DS, when ranked by the amount of 4-12 Hz power they contain. Each trace represents the difference of average PSDs for one channel in one patient; all 60 cortical channels are shown. In nearly all channels, power peaks at 6 Hz. H) Average time-frequency plot across subjects time locked to spindle starts from -1.25 to -0.75s prior to DS troughs. Downstates can be seen at -.25s before these spindle starts, marked by the pink arrow. Note that these spindle-preceding DS also precede the theta bursts, and the DS shown in panels A and D by about 1s, as shown by the two negative deflections in the superimposed gray waveform (color scale indicates ± 1dB, scale bar indicates 100 µV).

### Experimental Design and Statistical Analysis

Linear mixed effects models with subject specified as random effect were implemented in R to estimate descriptive statistics such as overall frequency, duration, and rate of occurrence, as well as for testing differences between frontal and parieto-occipital electrodes.

## Results

### Identifying short theta bursts prior to DSs in the cortex

Visual inspection of average spectrograms ± 2s relative to all DS troughs in intracranial recordings showed increases within the 5-10 Hz range approximately 250ms before the negative peak (Fig. 2 A,D, cyan arrows). However, inferring oscillatory activity from such representations can be misleading (Jones 2016), because the sharp decline in the pre-trough part of the DS could contain non-oscillatory power in the theta band (Cox et al. 2014). To confirm this increase was associated with an oscillatory component and not just an artifact of the ensuing DS LFP waveform, we looked for the presence of oscillations in the average of the original LFP time-locked to a filtered theta peak (5-8 Hz). We implemented this by sorting DSs according to previous theta power and averaging across events within a quartile, time-locked to the first band passed theta peak preceding the DS trough. This unfiltered average revealed clear oscillatory activity within the theta range, showing two to three peaks across the majority of channels for the top quartile and often absent in the lowest (Fig. 2B,E). Theta oscillations were also apparent at the level of single DSs selected from the top quartile (Fig. 2 B and E insets, cyan arrows).

To further characterize this observed intracranially-recorded theta oscillation during NREM sleep, we applied a theta detector to each channel (see Methods). Descriptive statistics for detected theta events are shown per subject in Table 2, and the overall estimates for frequency, duration, rate of occurrence are as follows (mean ± std. dev.): 6.33 ± 0.45 Hz, 672 ± 28 ms, 1.25 ± .39 /min. Interestingly, frontal channels exhibited lower overall frequency compared with parieto-occipital channels, with estimated frequencies of 6.26 and 6.5 respectively (t = -5.6, p = 2.14e-08, mixed effects model with subject as random effect). As expected, frontal channels also showed lower overall spindle frequency compared with parieto-occipital (t= -3.14, p=.002), with estimated overall frequencies of 12.26 and 12.65 Hz. To further verify that the pre-DS oscillations were the result of theta and not slower spindles, for each channel, we sorted DSs by power in 4-12 Hz 500ms prior to the DS trough. This range includes both theta and slow spindle frequencies, and thus would detect either. We then calculated the average periodogram across all DSs, per quartile. The difference of the top and bottom quartiles reveals a center of frequency at 6Hz across channels (2G), indicating that the pre-DS oscillations are theta, rather than slow spindles.

**Table 1.** Demographic and clinical information

| Patient | Sex | Age | Handedness | Clinical Diagnosis | Pathological Diagnosis | Imaging | Focus |
| --- | --- | --- | --- | --- | --- | --- | --- |
| 1 | M | 35 | - | Right temporal lobe epilepsy | No pathology obtained | Normal | Temporo-Parieto-Occipital Junction |
| 2 | F | 37 | R | Temporal lobe epilepsy | No pathology obtained | Normal | Hippocampus |
| 3 | M | 45 | L | CPS; bitemporal | No pathology obtained | - | Left and right mesial temporal lobes |
| 4 | F | 45 | R | CPS; multifocal | Multifocal: temporal, parietal, occipital | - | Temporal |
| 5 | F | 65 | R | CPS; temporal lobe epilepsy with two foci: left mesial temporal structures and right subfrontal region | No pathology obtained | - | Right subfrontal and anterior temporal |
| 6 | F | 32 | R | Left temporal lobe epilepsy | Non-specific gliosis | Normal | Hippocampus, entorhinal cortex, amygdala |
| 7 | F | 23 | R | Right temporal lobe epilepsy | Type I focal dysplasia | Normal | Hippocampus, entorhinal cortex, anterior insula |
| 8 | F | 28 | R | Right temporal occipital epilepsy | Type II focal dysplasia | Focal cortical dysplasia in right fusiform | Fusiform gyrus, entorhinal cortex |
| 9 | F | 50 | R | Right temporal occipital epilepsy | No pathology obtained | Normal | Right fusiform gyrus |
| 10 | F | 27 | R | Right temporal lobe epilepsy | No pathology obtained | Right hippocampal sclerosis | Hippocampus, entorhinal cortex, amygdala |

**Table 2:**
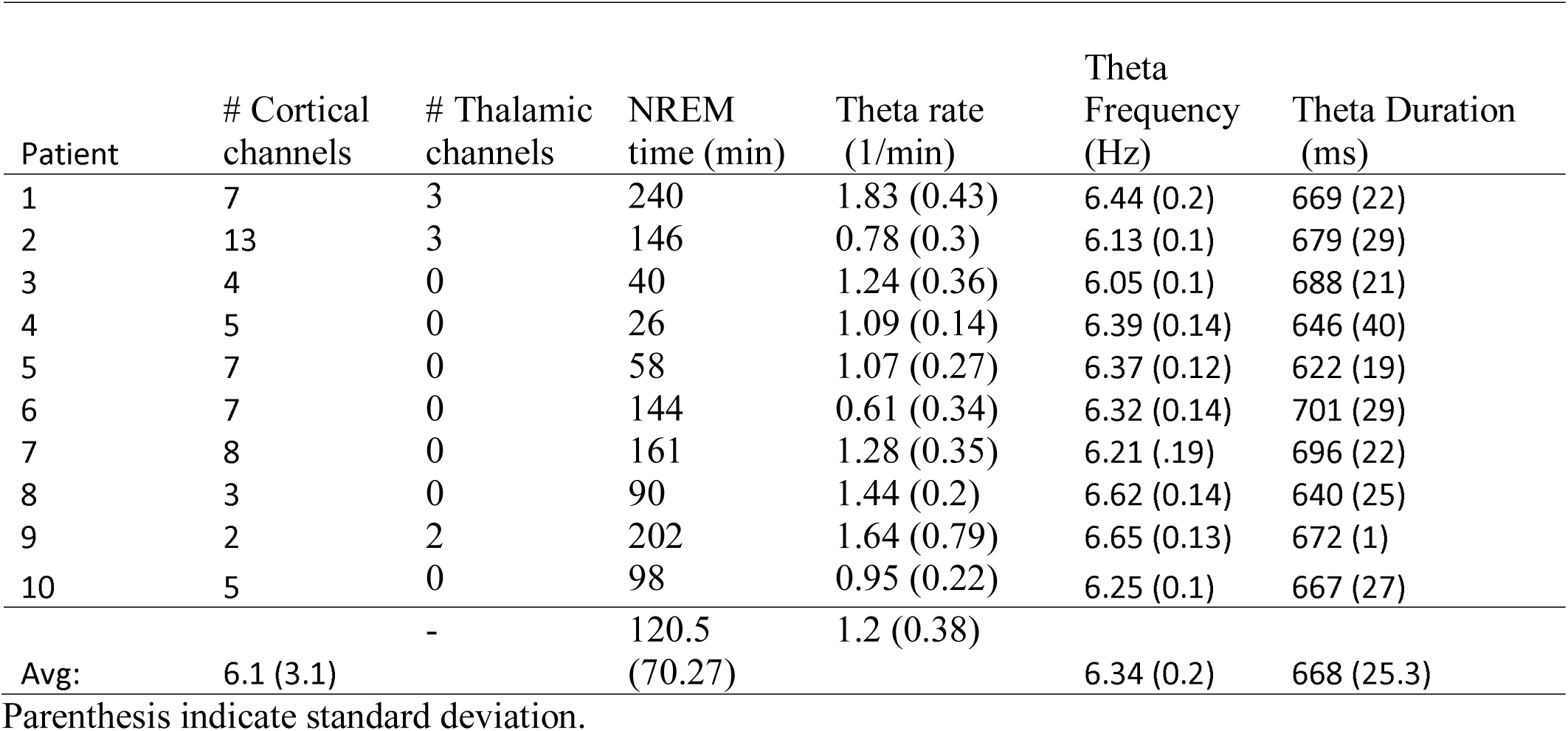
Theta burst characteristics in thalamus and cortex

In Figures 2C and 2F, we superimpose all detected theta traces, unfiltered and locked to the deepest trough in the theta event for two example bipolar SEEG channels. Some channels exhibited a downward trend in the average (Fig. 2F), suggesting a DS tends to follow the deepest trough, whereas others did not (Fig. 2C). This difference illustrates that while some detected theta events were associated with DSs, others were not, and that this varied within and between channels. We also observed from *post-hoc* analyses that some channels exhibited larger amplitude and more prolonged theta oscillations in the raw LFP for N2 versus N3 DSs (Fig. 3A). This was corroborated with a greater number of peaks in detected theta events on average per channel for N2 compared with N3 (t = -4.33, p = 1.49e-05, mixed effects model with subject as random effect and channel as nested random effect). Additionally, we found there was significantly greater theta power prior (ࢤ500ms to 0) to the DS trough for N2 compared to N3 DSs (Fig. 3B; t = -3.13, p = .002, mixed effects model). However, the rate of detected theta event occurrence was not different between N2 and N3 (t = -1.2, p = .23). That is, although the number of events did not differ between N2 and N3, theta bursts were longer and larger in N2.

**Figure 3.**
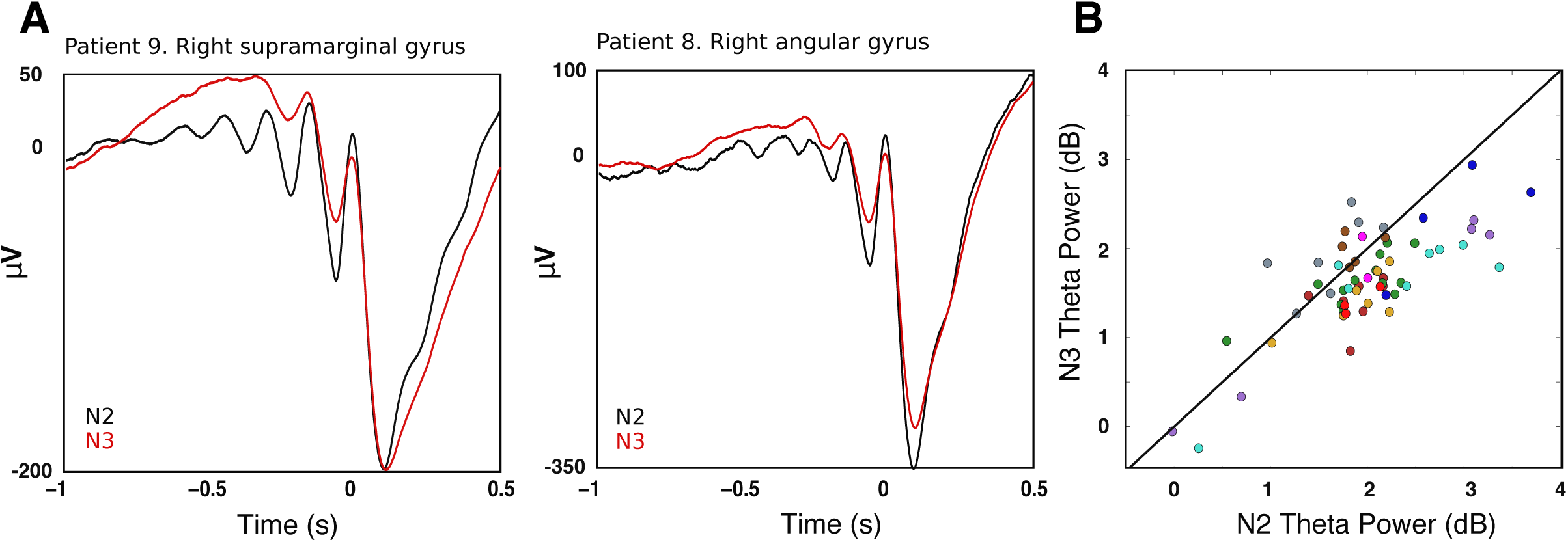
Comparing N2 and N3 theta bursts. A) Theta oscillations prior to DS troughs were more pronounced for N2 than N3 DSs. Average waveforms of the DS with the top quartile of prior theta power are shown from two patients within N2 in black (n=677 in pt 9 and 174 in pt 8) and N3 in red (n=913 in pt 9 and 635 in pt 8). Averages are locked to the theta peak just preceding the DS trough. B) Power within theta (5-8Hz) range -500ms to 0 relative to DS trough for each channel (channels from a given patient have the same color). Most dots are below the diagonal, indicating theta power was greater for N2 than N3 (t = -3.13, p = .002, mixed effects model with subject as random effect and channel as nested random effect).

### Theta and spindles show different temporal relationships with DSs

Since slow oscillations during sleep are known to group higher-frequency rhythms, we investigated how often our detected theta bursts (TBs) occurred in relation to DSs recorded intracranially. To quantify this, we created event-related histograms for each bipolar channel by relating the start of TBs to DS troughs. For example, a recording from Patient 1 in the right middle frontal gyrus shows that when TBs are detected around DSs, they are more likely to start approximately 300ms prior to the DS trough (Fig. 4A). At the same cortical location, the likelihood of slow spindles (Fig. 4B), fast spindles (Fig. 4C), or all spindles (Fig. 4D) starting between -500ms to 0 is greatly reduced compared to after the DS trough.

**Figure 4.**
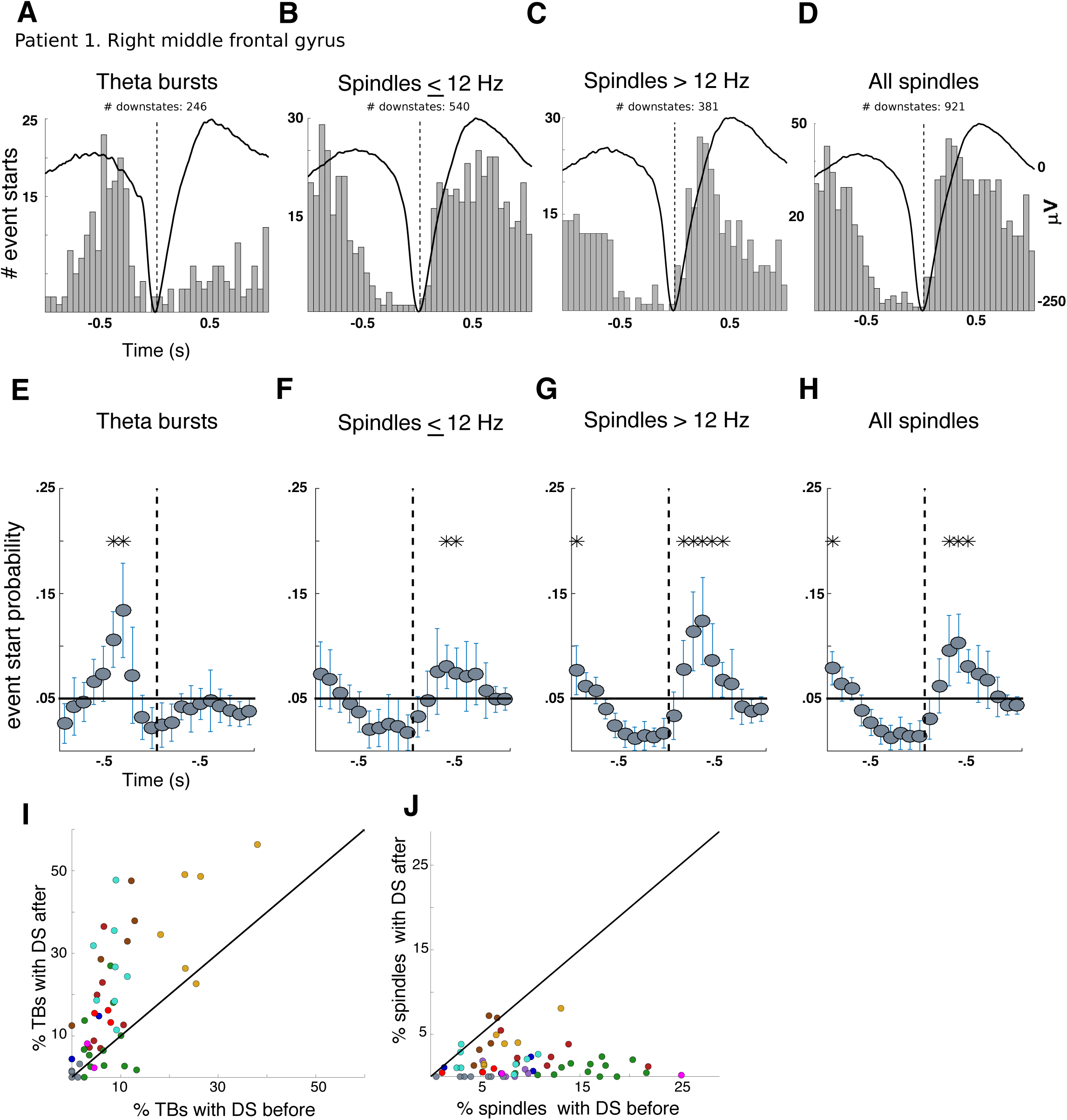
Cortical theta precedes DSs, both slow and fast spindles follow. A-D) Event-related histograms showing the timing of the start of theta events (A), spindles ≤ 12 Hz (B), spindles >12 Hz (C), and all spindles (D) relative to the DS trough at a single bipolar channel in the right middle frontal gyrus of Patient1 during both N2 and N3 sleep. E-H) Histograms from all channels, normalized by the total number of counts ±1s and pooled in 100ms bins. The black line indicates chance level, and probability estimates per bin across channels were calculated using linear mixed effects models. Stars indicate bins where events occur significantly more than chance (Bonferroni adjusted, p<0.05); error bars indicate 95% CIs. I) Scatterplot, for each channel, of the proportion of theta bursts with the DS within 500ms after (y-axis) versus the proportion with the DS within 500ms before (x-axis); J) Same as I but for spindles. All cortical channels with at least 50 events occurring within ±500ms of the DS trough were included. Most channels (dots) in I are above the diagonal, indicating that the DS trough usually occurs after TB onset; in contrast, most channels in J are below the diagonal, indicating that the DS trough usually occurs before spindle onsets.

Of 60 cortical recordings, 35 channels from 7 patients had at least 20 TBs within ± 500ms of the DS trough, none of which were significantly more likely to start after the DS trough. However, 14 of the 35 channels from 5 patients were more likely to have TBs start before the DS trough (Binomial test, Bonferroni adjusted p<0.05). The proportion of TBs occurring prior to the DS trough for these channels was not significantly different between frontal and parieto-occipital regions (t=1.1, p=0.27).

Previous work (Mölle et al. 2011; Klinzing et al. 2016) asserts that spindles have different temporal relationships with DSs depending on spindle frequency; with slower spindles (≤ 12 Hz) occurring on the up to down transition, and faster spindles (>12 Hz) occurring on the down to up transition. In order to test this hypothesis, we selected from the 60 cortical recordings, the 35 channels from 7 patients which had at least 20 slow spindles within ± 500ms of the DS trough. None of these channels showed slower spindles which were significantly more likely to start before the DS trough. However, slower spindles recorded by 20 of the 35 channels from 6 patients were more likely to start *after* the DS trough (Bonferroni adjusted p<0.05). If all spindles were grouped together, those recorded by 39/49 channels from 10 patients were significantly more likely to start after a DS trough (Bonferroni adjusted p<0.05). Neither slower spindles (t=-.44,p=0.66) nor all spindles (t=-1.25, p=0.21) showed significant differences between frontal and parieto-occipital regions in the proportion of spindle events occurring after DS troughs.

Normalizing each event-related histogram per channel by the total number of counts in the ±1s time window and pooling histograms across all cortical bipolar channels from 10 subjects revealed similar results (Fig. 4E-H): TBs initiate before DS troughs and spindles—both slow and fast--initiate after. Significance was assessed by Bonferroni correction (p<0.05) across time bins within each event type. Time bins with significant likelihood of TBs starting were centered around -450 and -350 ms before the DS trough (Fig. 4E). In contrast, slower spindles were significantly more likely to start 350-450 ms after the DS trough (Fig. 4F), and faster spindles were significantly likely to start between bins centered 150-550ms after the DS trough (Fig. 4G). This confirms that spindles are more likely to start during the transition from down to up states, while TBs start just before the transition to a DS.

Some time bins (Fig. 4B-D,G & H) showed increased likelihood of spindles starting from around -1.25s to -0.75s prior to DS troughs. To see if this increase was due to a previous DS, we identified the ‘early’ spindles in question as those that began around -1s before down state troughs (between -1.25s and -.75s relative to the DS minimum). Then we generated a time-frequency plot for the activity surrounding these spindles for each channel, and then averaged them across the 33 channels in six subjects which had at least 50 such spindles (Fig 2H). A downstate is clearly present in the average time-frequency plot, peaking about .25s prior to the onset of these ‘early’ spindles, corresponding to the DS at about -1s in plots 2AD. The same plots show another downstate at about 1s following ‘early’ spindle onset; this is the downstate that was used to identify the ‘early’ spindles in the first place, corresponding to the DS at 0.0s in plots 2AD. Notably, the ‘early’ spindle power increase precedes the theta band increase in this plot, as well as those triggered on the main downstate (Figs. 2A and 2D). In summary, we demonstrate that there is a sequence of sleep grapho-elements, typically theta-downstate-spindle, but sometimes downstate1-spindle1-theta-downstate2-spindle2. The second, longer sequence is expected given that, especially in stage N3, downstates are well-known to occur rhythmically at about 1Hz, comprising the slow oscillation.

Next we asked, how often do TBs or spindles occur around DSs? Only channels that had at least 50 occurrences of each event type (theta or spindle) were included (N = 9 patients, 49 cortical channels). Downstate masks were padded with 100ms on either side of the DS trough, and spindle and theta event masks marked by estimated start times. We found on average across subjects, 24% of detected TBs began within ± 500ms of a DS trough, with 7.7% of TBs occurring after a DS and 17.8% occurring before (Fig. 4J). In contrast, 10% of detected spindles began within ± 500ms of a DS trough, with 8.7% of spindles occurring after a DS and 1.7% occurring before (Fig. 4K). There was a significantly greater proportion of TBs that fell within ± 500ms of a DS trough than spindles (paired t-test, p=2.4e-05). There was substantial inter-subject variability for both events, especially for TBs. Despite this variability, most subjects exhibited similar temporal relationships with DSs.

### Relation of theta bursts and spindles recorded in scalp EEG to downstates

In order to ensure that that our findings generalize to non-patient populations and to contextualize our results within the framework of more commonly recorded non-invasive measures, we detected events recorded using scalp EEG in three healthy human participants and re-calculated event-related histograms, as in Fig 2 E-H. Replicating the SEEG results reported above, normalized peri-downstate histograms revealed divergent patterns for theta bursts and spindles recorded at the scalp (Fig 5): TBs initiate before DS troughs, whereas spindles begin afterwards. Significance was assessed by Bonferroni correction (p<0.05) across time bins within each event type. TBs had a significant likelihood of starting 500-300 ms before the DS trough (Fig. 5A). In contrast, slower and faster spindles were significantly more likely to start during time bins centered on 450ms and 350ms, respectively, after the DS trough (Fig. 5B & C). These results are in concordance with our conclusions from SEEG: spindles are more likely to start during the transition from down to up states, while TBs start just before the transition to a DS.

**Figure 5.**
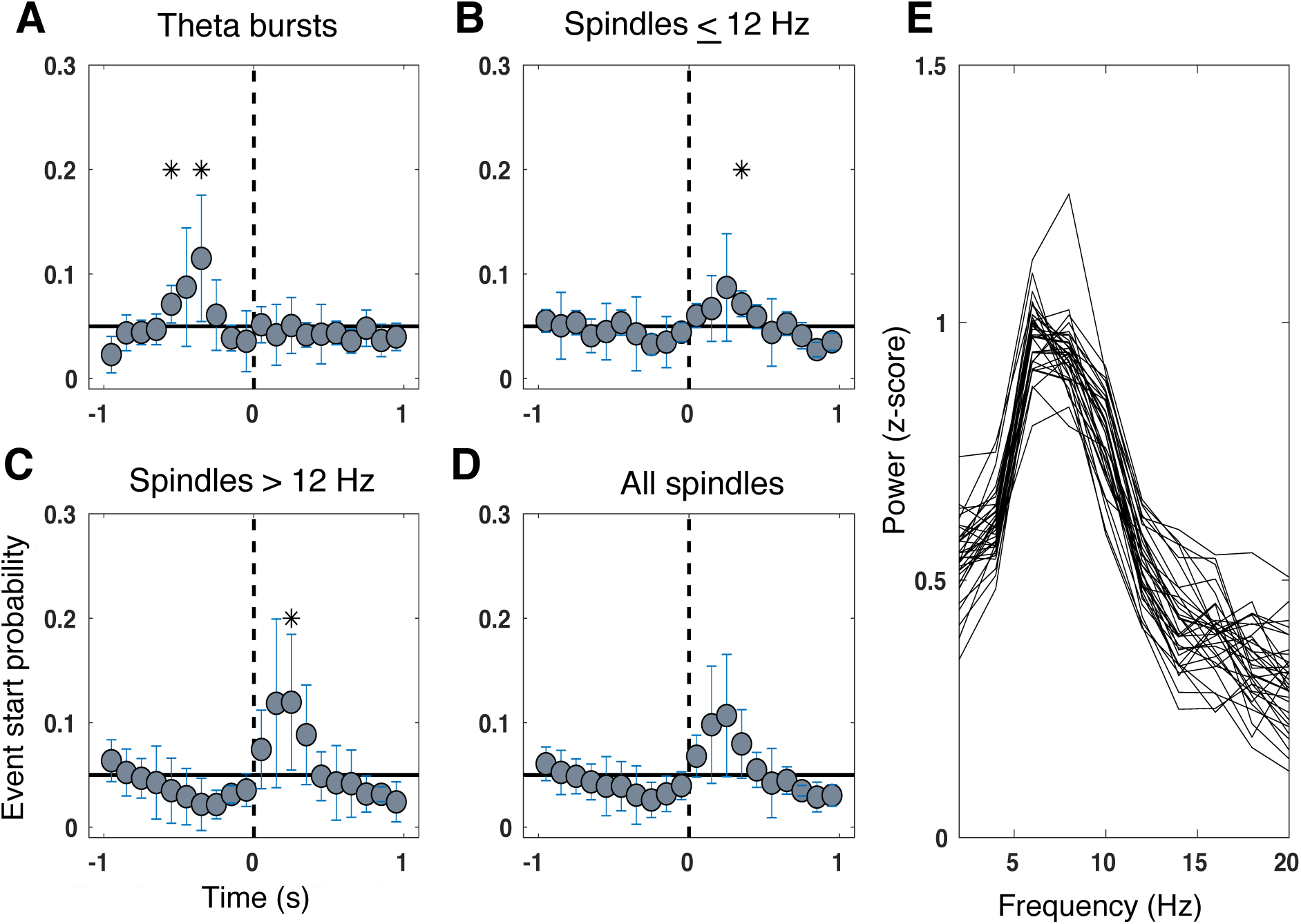
Scalp EEG theta precedes DSs, spindles follow. As in Fig. 4E-H, DS-locked event-related histograms from all scalp EEG channels, normalized by the total number of counts ±1s and pooled in 100ms bins are plotted for the start times of: A) theta bursts, B) spindles ≤ 12Hz, C) spindles > 12Hz, and D) all spindles. The black line indicates chance level, and probability estimates per bin across channels were calculated using linear mixed effects models. Stars indicate bins where events occur significantly more than chance (Bonferroni adjusted, p<0.05); error bars indicate 95% CIs. E) Pre-downstate spectra. Plotted are the difference of the first and fourth quartile average power spectral densities of 500 ms pre-DS, when ranked by the amount of 4-12 Hz power they contain. Each trace represents the difference of PSDs for one channel, averaged across all subjects. Only channels common to all subjects are included.

### Relating thalamic theta with cortical theta

Three patients also had SEEG electrodes which recorded from the thalamus, and we recently characterized the coordination of NREM DS and spindles between cortex and thalamus in these patients (Mak-McCully et al. 2017). Here, we investigated whether the thalamus also exhibits similar TBs to the cortex, and if so, how they relate to cortical TBs.

Both time-frequency representations locked to DSs (Fig. 6A, example channel) and averaging the raw LFP for the top and bottom quartiles of prior DS theta power (Fig. 6B, example channel) suggest there are similar TBs before DSs in the thalamus. These oscillations could also be seen at the level of single DSs (Fig. 6B inset), and often revealed a downward slope in the average LFP of detected TBs (Fig. 6C), similar to some cortical channels. Compared with cortical channels from the same subjects (shown in Table 2), thalamic channels showed no difference in the rate of theta occurrence (mean ± std. dev. for thalamus= 1.07 ± 0.66; for cortex= 1.39 ± 0.62; t=1.4,p=.16), average frequency (mean ± std. dev. for thalamus= 6.25 ± 0.13; for cortex= 6.33 ± 0.22; t=1.12,p=.26), or duration (mean ± std. dev. for thalamus= 661 ± 17 ms; for cortex= 675 ± 25 ms; t=1.43,p=0.15) in detected events. This is in contrast with DSs, which showed greater rates of occurrence in the cortex, and spindles, which occurred more frequently in the thalamus, as shown in our previous study (Mak-McCully et al. 2017).

**Figure 6.**
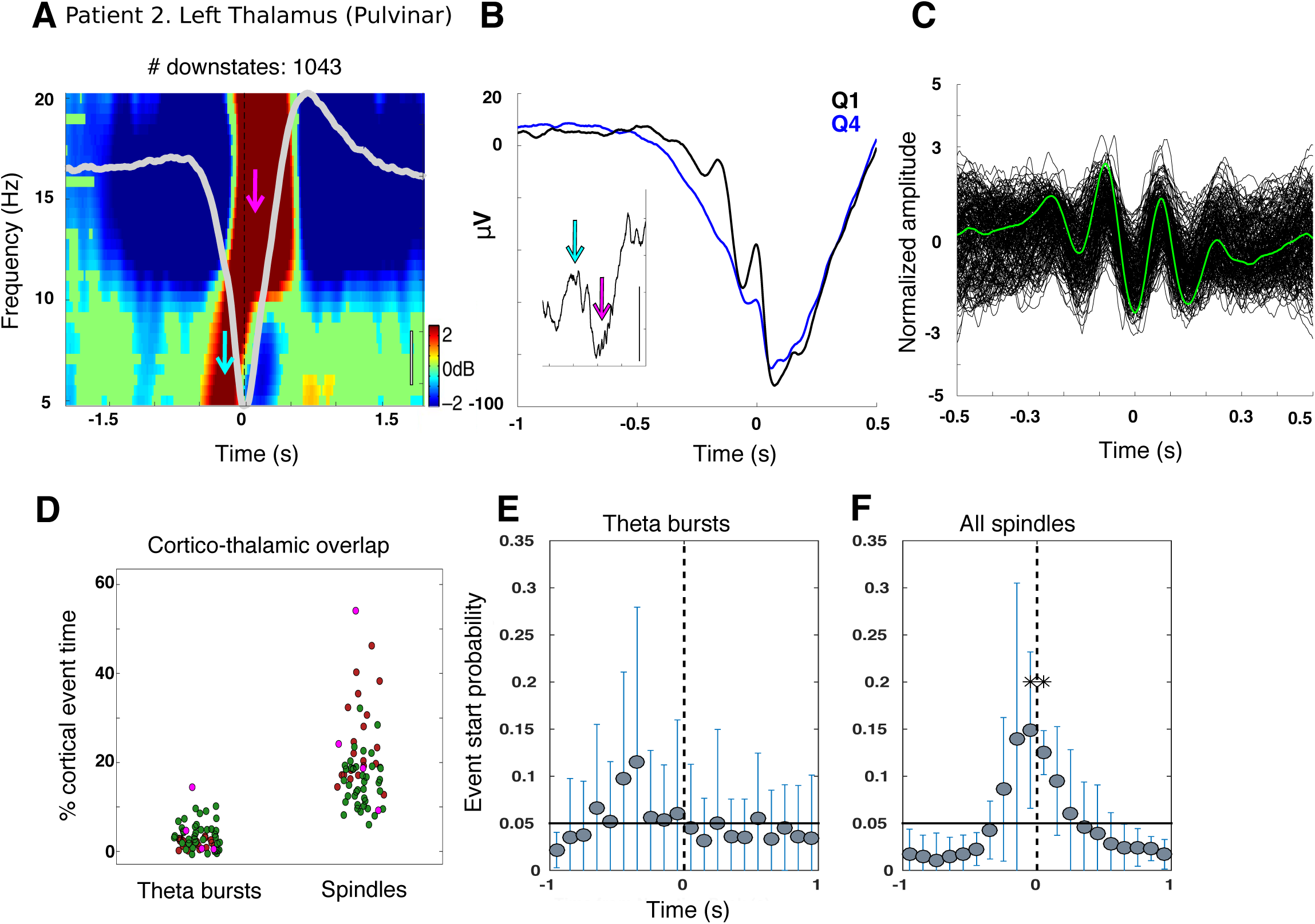
Thalamic theta during NREM sleep. A) Example time-frequency representations locked to DS troughs for a bipolar thalamic channel from Patient 2 showing increased power in the theta range prior to the trough (cyan arrow) and in the spindle range just after the trough (pink arrow). The average DS is overlaid in light gray and black scale bar indicates 20µV.See Mak-Mcully et al 2017 Fig 2B for time-frequency plots extending to 100Hz. B) prior theta power is not solely driven by DS waveform shape, but displays oscillatory activity in the theta range after averaging the raw LFP locked to the first theta peak preceding DS the trough (as in Fig. 2B & E). The inset displays a single trace from the top quartile for ± 1s around DS trough, scale bar indicates 100µV; cyan and pink arrows pick out individual theta and spindle events. C) All detected theta events locked to the deepest trough for the same channel as A and B. D) The proportion of cortical and thalamic overlap for theta bursts and spindles. Each dot is a different channel; dots from the same patient share the same color. Y-axis indicates the proportion of time a given cortical channel has a theta burst (spindle) overlapping with a thalamic theta burst (spindle) after subtracting the amount of overlap expected by chance. Spindles show a greater degree of corticothalamic overlap compared with theta bursts (t=14.2, p=1.2e-45; subjects as random effects, corticothalamic pairs as nested random effect).

We found that cortical TBs only slightly overlapped with thalamic TBs after correcting for the overlap expected by chance (average overlap per patient = 1.9,3.7, and 5.1 % above chance), but this overlap was nonetheless significant (t=4.04, p= 5.3e-05; subject as random effect, corticothalamic pair as nested random effect). In contrast, cortical spindles in most sites overlapped strongly with thalamic spindles over the proportion expected by chance (average overlap per patient = 24.2, 15.6, 26.6% above chance), and again this was highly significant (t=6.22, p=5e-10 Fig. 6D). This greater corticothalamic overlap for spindles compared to TBs is significant (t=14.2, p=1.2e-45).

We also examined the co-occurrence of thalamic and cortical theta events using their joint occurrence histograms. Only 5/64 corticothalamic pairs had at least twenty thalamic TBs starting within ± 500ms of the start of cortical TBs, and none of these 5 pairs had histograms with a significant difference between leading versus lagging peaks as assessed with a binomial test (p<0.05, Bonferonni corrected). This is in contrast to spindles, which showed significant ordering effects with the thalamus preceding the cortex, and DSs, which usually started in the cortex prior to thalamus (Mak-McCully et al. 2017). These findings provide support that our detected theta events are separate entities from DSs and spindles, as they appear to have distinct corticothalamic dynamics.

When relating thalamic TBs to thalamic DSs, 6 of 8 thalamic channels from three patients had at least 20 TBs within ± 500ms of the DS trough. Of these 6, two channels from two patients were more likely to have TBs start before the DS trough (Binomial test, Bonferroni adjusted p<0.05); no channels were more likely to have TBs start after the trough. When pooled across patients, TBs were not more likely to start in any time bin over chance, relative to DS troughs (Fig. 6E). In contrast, peak occurrence of thalamic spindle onset was at the thalamic downstate peak (Fig. 6F), as has been previously reported (Mak-McCully et al. 2017).

### Theta and spindles differ in high frequency coupling

To determine if the TBs prior to DSs modulated HG power, we took the top quartile of DSs with prior theta power (5-8Hz, -500ms to 0 negative peak), and averaged both the HGP (60-100Hz) and LFP locked to the band passed theta peak first preceding the trough in the bipolar cortical channels. This revealed clear oscillations within the theta range in the HGP (Fig. 7A, example channel). We performed a similar analysis for spindles in the same channels, only with DSs sorted by spindle power 0 to 500ms after DSs (Fig. 7B), and averaging time-locked to the first post-DS filtered peak in 10-16 Hz. To quantify the extent of this modulation, we employed traditional cross-frequency coupling metrics at the beginning (first two cycles) of detected spindle and TBs (See Materials and Methods).

**Figure 7.**
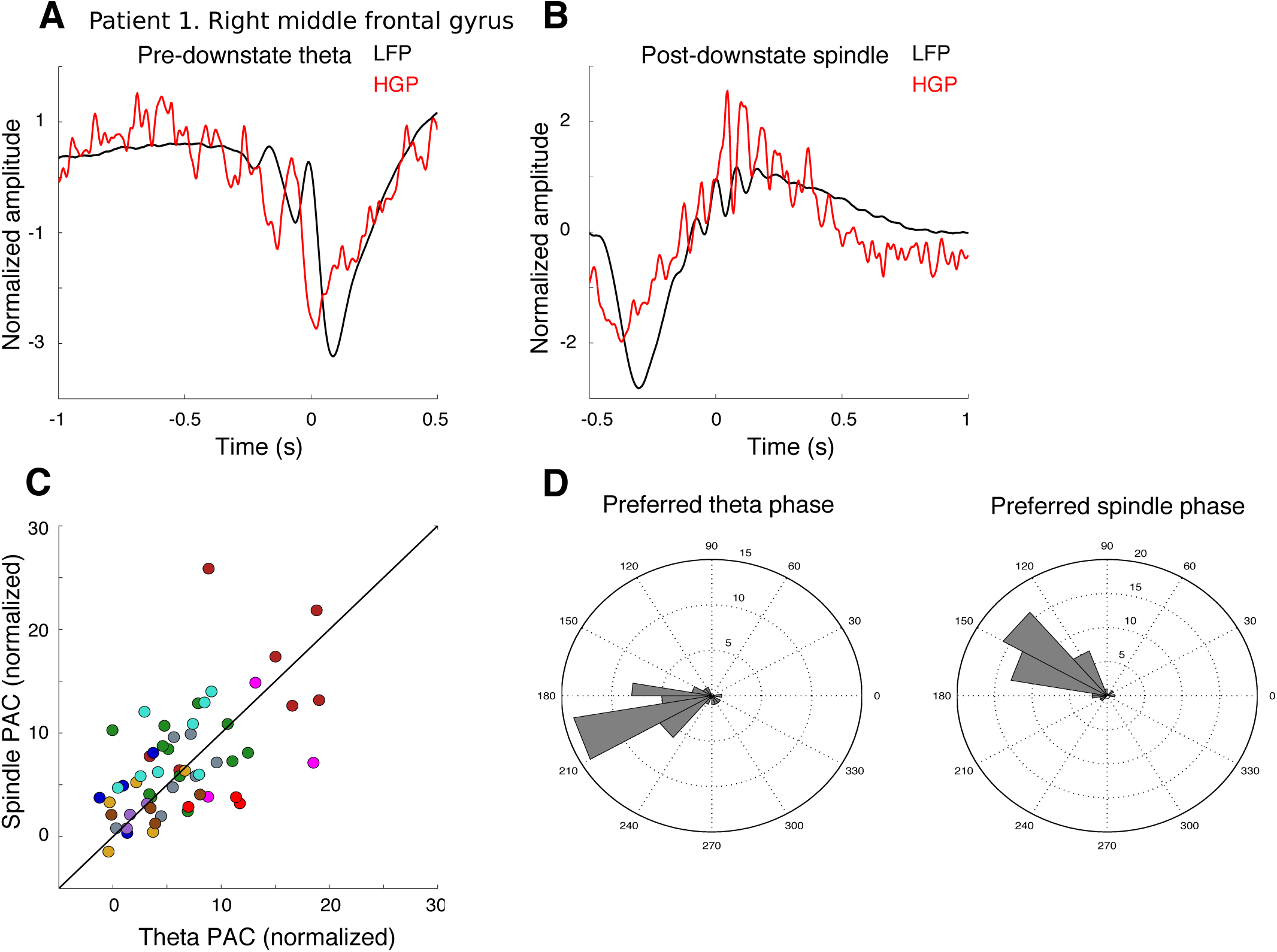
Theta and spindles couple to high gamma power differently. A,B) Patient 1, right middle frontal gyrus. A) Average of the top quartile of DSs (ranked by pre-DS theta power) locked to the first prior theta peak (black), as well as the average of HGP (60-100Hz) for the same events (red). B) Average of the top quartile of DSs (ranked by post-DS spindle power) locked to the first filtered spindle peak following the DS trough (black), as well as the average of the HGP. C) Magnitude of phase-amplitude coupling (PAC) between theta phase and HG (x-axis), versus spindle phase and HC (y-axis), for all bipolar channels. Overall, a similar level of coupling is observed for theta bursts vs spindles. D) Preferred phase of either theta or spindle events for each channel that had significant PAC. Radial scale is number of channels. The preferred phase is consistent for each type of wave and differs significantly between theta bursts and spindles (parametric Watson-Williams, F=26.5,p=1.6e-6).

We found that for TBs, 42 cortical channels from 10 subjects showed significant PAC-Z (p<0.002, uncorrected for multiple comparisons), and 45 channels from 10 subjects were significant for spindle events. There was a strong correlation between PAC-Z measures from the two event types across channels (Fig. 7C; spearman rho =0.59, 10 subjects, 57 channels), and no difference in amplitude of PAC between theta and spindle events (p=0.15; N=10, 57 channels; nested random effects of channel in subject). However, there was difference in the preferred phase for HG power for TB (at ~200°) and for spindles (at ~140°; Fig. 7D). This difference was highly significant (parametric Watson-Williams, F=26.5,p=1.6e-6).

## Discussion

Loomis’ original recordings of sleep EEG (1939) commented on sleep spindles following K-Complexes (i.e., DS:(Cash et al. 2009)). This coupling was later quantified (Mölle et al. 2002), and then extended to assert that while ‘fast spindles’ occur after downstates, ‘slow spindles’ occur before (Mölle et al. 2011; Klinzing et al. 2016; Yordanova et al. 2017). Like these studies, we found contrasting patterns of oscillatory activity before versus after DS. However, unlike these studies we found that both slow and fast spindles occurred post-DS in direct cortical recordings from both frontal and occipito-parietal sites (Fig. 4), as well as scalp EEG (Fig. 5). Rather than slow spindles, we found that short theta bursts (TB) precede DS, whether in cortical or scalp recordings. Increased spectral power preceding DS is mainly within the theta band, centered at ~6Hz (Fig. 2G, 5E), but extends into the spindle range. Consequently, if this activity is band-passed in the spindle range, low frequency ‘spindles’ could be detected despite the center frequency of unfiltered recordings being in the theta band.

Previous scalp studies have reported increased theta power prior to DS troughs (Cox et al. 2014; Klinzing et al. 2016). We confirmed that these were true oscillations by averaging the raw LFP locked to a theta peak prior to the DS (Fig. 2B & 2E), and by requiring that each potential theta burst contain at least three peaks. Theta bursts thus consist of multiple 4-8 Hz waves, with a mean duration of ~670ms, shorter than most spindles. They occur at about the same rate in both N2 and N3 but have fewer cycles and lower amplitude in N3. The density of TB is about 7 fold less than spindles, and unlike spindles (Mak-McCully et al. 2017), TB density does not differ between cortex and thalamus. In addition, while spindles are tightly coupled between thalamus and cortex, no significant relationship can be observed for TB. Finally, cortical TB have a significantly different phase relation to HG than spindles, indicating distinct generators (Fig. 7D). Thus, TB are distinguished from spindles in their internal frequency, duration, density, position relative to the DS, lack of overlap or consistent sequencing between thalamus and cortex, and distinct HG phase preferences.

If slow versus fast spindles do not differ in their relation to the DS, then is it still tenable to claim that they represent distinct neurophysiological phenomena rather than variations within a continuum? Our study confirms the slightly but significantly higher average frequency of spindles in parietal versus frontal cortex, consistent with previous EEG and MEG (Dehghani et al. 2011), and intracranial recordings (Andrillon et al. 2011; Peter-Derex et al. 2012; Piantoni et al. 2017). The individual waves in EEG and MEG spindle bursts also vary in frequency, with later waves also ~1Hz slower on average (Dehghani et al. 2011). In SEEG, within a given cortical location, and often within the same spindle, spindle waves with frequencies both above and below the fast/slow division are typically observed. Two kinds of spindles can be identified in laminar recordings, involving mainly upper or middle layers, respectively (Hagler et al. 2018). The average frequency of upper versus middle channel spindles does not differ significantly, and both include both slow and fast spindle waves. In all of these circumstances, fast and slow spindles occur in a continuum rather than a dichotomy.

Within this framework it is not clear how to explain how, in some subjects, two peaks in the spindle spectrogram can be discerned at the scalp (Cox et al. 2017). The cortical origin of scalp EEG spindles is not yet well understood due to the lack of detailed information regarding the amplitude, density, synchrony, phase and orientation of the generating cortical patches. It is thus theoretically possible that the slower spindles reported at the scalp are from a location where we did not record. However, the parietal and frontal cortices where we recorded have been proposed to be the generators of fast and slow scalp spindles (Mölle et al. 2011; Klinzing et al. 2016), and generate spindle band activity most related to scalp EEG spindles (Frauscher et al. 2015). Because the inverse problem is ill-posed, it is possible to model the scalp EEG spindle distribution as due to either anatomically distinct generators, each with a single frequency, or distributed generators, each with a range of overlapping frequencies, changing slightly across areas. Our results clearly support the second model.

This supposed dichotomy between slow pre-DS spindles and fast post-DS spindles in humans has been homologized to the clear dichotomy in rodents between high voltage slow spindles (~8Hz) vs. low voltage fast spindles (~14Hz) (Timofeev & Chauvette 2013). Only the fast spindles are associated with memory replay and consolidation (Eschenko et al. 2006; Johnson et al. 2010), or DS (Johnson et al. 2010). The sharp waveforms and other epileptiform characteristics of slow rodent spindles (Polack 2006) suggest that they do not have an homology in healthy human recordings. Thus, it appears that fast rodent spindles correspond to both faster and slower spindles in humans.

The TB preceding DS in N2 comprise an augmenting oscillation between cortical excitation and inhibition as indexed by phase-locked high gamma, which is correlated with neuronal firing (Lachaux et al. 2012). The greatest HG decrease occurs at the final surface-negative TB trough, which coincides with the DS trough. Thus, in TB-DS sequences, the DS does not arise as a sudden decline from baseline, but as the culmination of an escalating TB oscillation, suggesting that the TB may play a role in helping to trigger the DS as its final cycle. This possibility receives some indirect support from the fact that they are generated by the same cortical layers in laminar recordings (Halgren et al. 2018; Csercsa et al. 2010) and thus may be engaging the same circuits. The hypothesis that theta waves may trigger DS in N2 provides a solution to a difficult problem: how do KCs arise? Current theories model DS onset as a response to the preceding upstate (Neske et al. 2016). This view describes the usual *in vitro* or anesthetized recordings where US arise from a flat depressed baseline, which is considered the DS (Lemieux et al. 2014). In contrast, in unanesthetized humans (Mak-McCully et al. 2015) and animals (Chauvette et al. 2011), DS appear as stereotyped events in a chronically active cortex during NREM sleep. Notably, laminar recordings in humans demonstrate that KCs are DS without a preceding US (Cash et al. 2009). We suggest here that the last positive peak of the TB may replace the US as the DS trigger. For example, the calcium influx associated with this cortical excitation during the final positive peak would trigger hyperpolarizing K+ currents (Cunningham et al. 2006), which may tip the circuit into the DS.

The consistent relation of TB to DS, and DS to spindles, may be related to the appearance of these waves in successive stages of sleep. The transition from quiet waking to N1 is marked by the replacement of alpha by theta waves. N2 then appears when KCs and sleep spindles appear. Simultaneous corticothalamic recordings in natural human sleep show that converging cortical DS precede thalamic DS, that thalamic spindles are tightly coupled to begin at the thalamic DS trough, and that thalamic spindles drive cortical (Mak-McCully et al. 2017). In this view, the N1 to N2 transition would occur when theta begins to trigger cortical DS (i.e., KCs), which in turn trigger successively thalamic DS, thalamic spindles, and cortical spindles. N2 transitions to N3 when DS and upstates recur rhythmically as the slow oscillation. Both TB (shown here) and spindles (Andrillon et al., 2011, Mak-McCully et al., 2017, Piantoni et al., 2017) continue during N3, but become abbreviated. We hypothesize that as the neuromodulatory state deepens, the US triggered by the DS becomes capable of triggering the following DS, resulting in less time for elaboration of the TB or spindles.

During NREM sleep, hippocampal cells replay events from the preceding waking period during sharp-wave ripples, which arrive at the cortex during the down to upstate transition, as the spindle is beginning (Maingret et al. 2016; Jiang et al. 2017). This conjunction of hippocampal input with cortical modulation is thought to underlie consolidation of cortical memory circuits (Diekelmann & Born 2010; Hanert et al. 2017; Niknazar et al. 2015; Mölle et al. 2011; Latchoumane et al. 2017). Our results suggest that TB may precede and could help initiate the downstate-ripple-spindle-upstate sequence. During waking, theta occurs during tasks requiring sustained processing (Kahana et al. 1999; Raghavachari et al. 2006), and may underlie prominent cognitive event-related potentials (Halgren et al. 2015; Cavanagh et al. 2011). If theta serves a similar function during NREM, then TB may organize the gathering of cortical information prior to the DS, and consequently help select related hippocampal traces to be activated and sent back to the cortex for integration with currently active neurons during the upstate and spindle.

## Notes

**Acknowledgements:** This work was supported by the National Institutes of Health Grants R01-MH-099645 and R01-EB-009282, the U.S. Office of Naval Research Grant N00014-13-1-0672, the National Science Foundation Graduate Research Fellowships Program, and the National Institute of Mental Health T32 Cognitive Neuroscience Training Grant. We thank Nima Dehghani for EEG data, Donald Hagler for spindle detection scripts, Fabrice Bartolomei for access to data and analysis input, Catherine Liegeois-Chauvel for research access, Jean Regis for electrode localization for the Marseille patient.

